## Supplementary Data for "Paradoxical role of AT-rich interactive domain 1A in restraining pancreatic carcinogenesis"

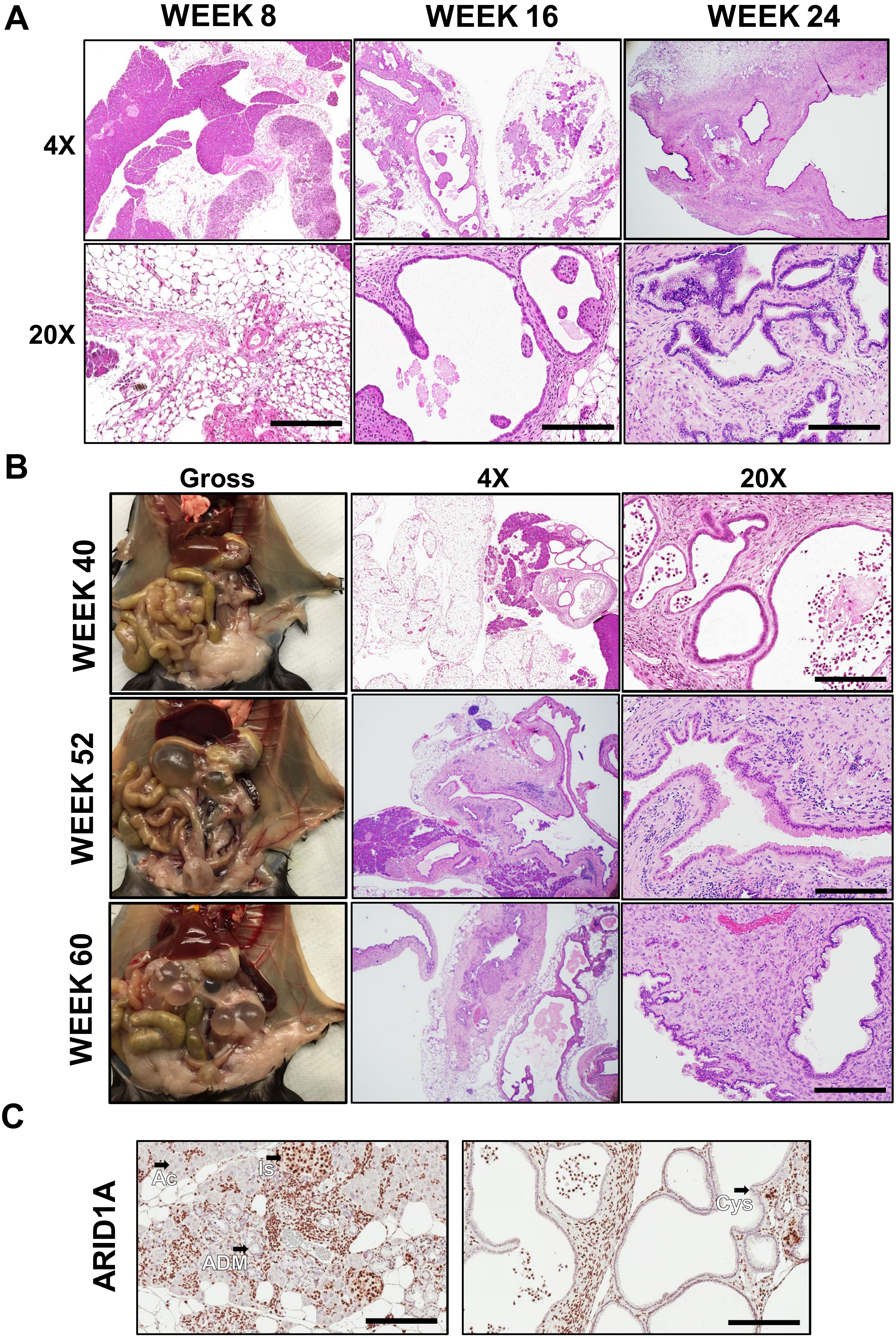


SUPPLEMENTAL FIGURE S1

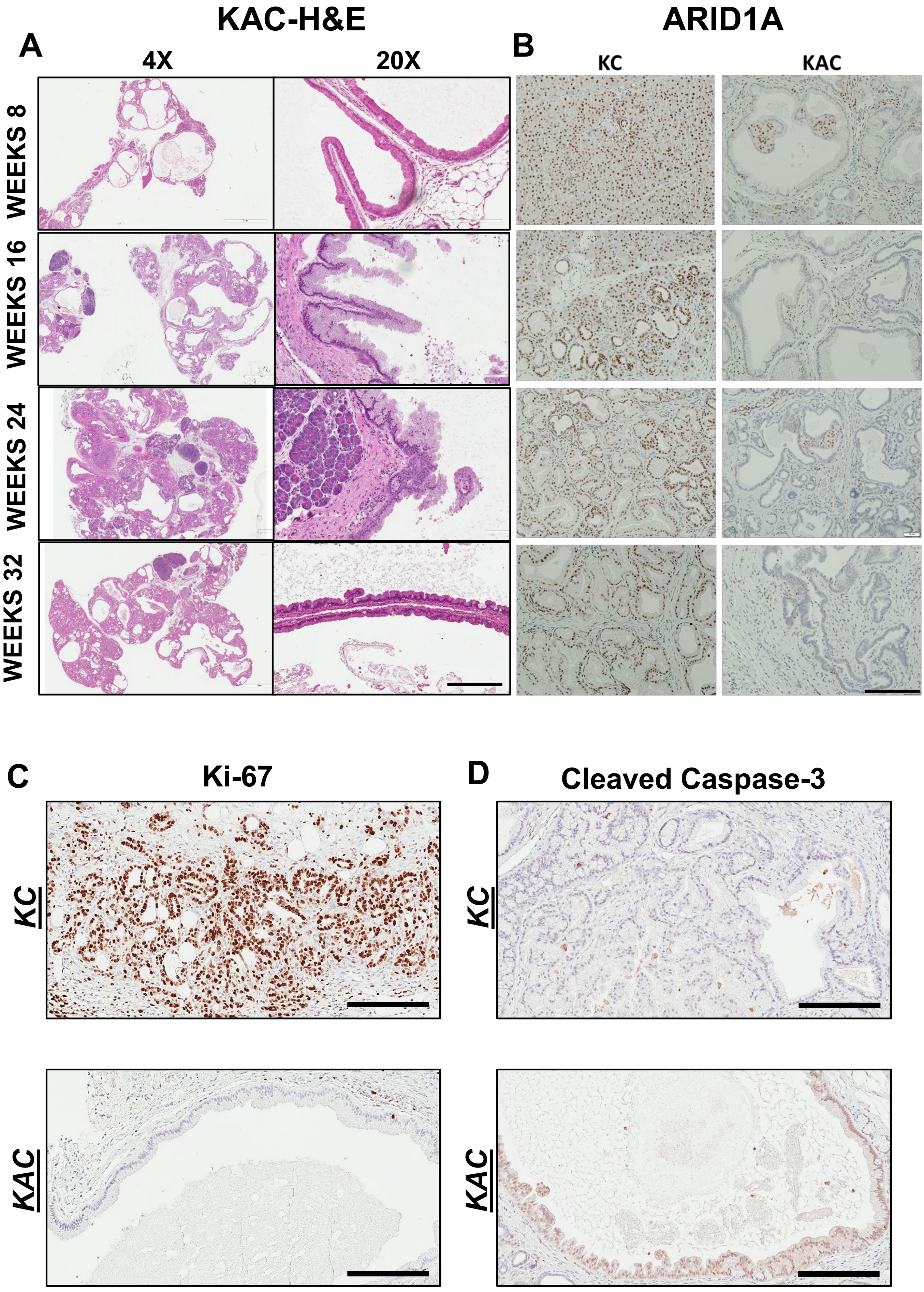


SUPPLEMENTAL FIGURE S2

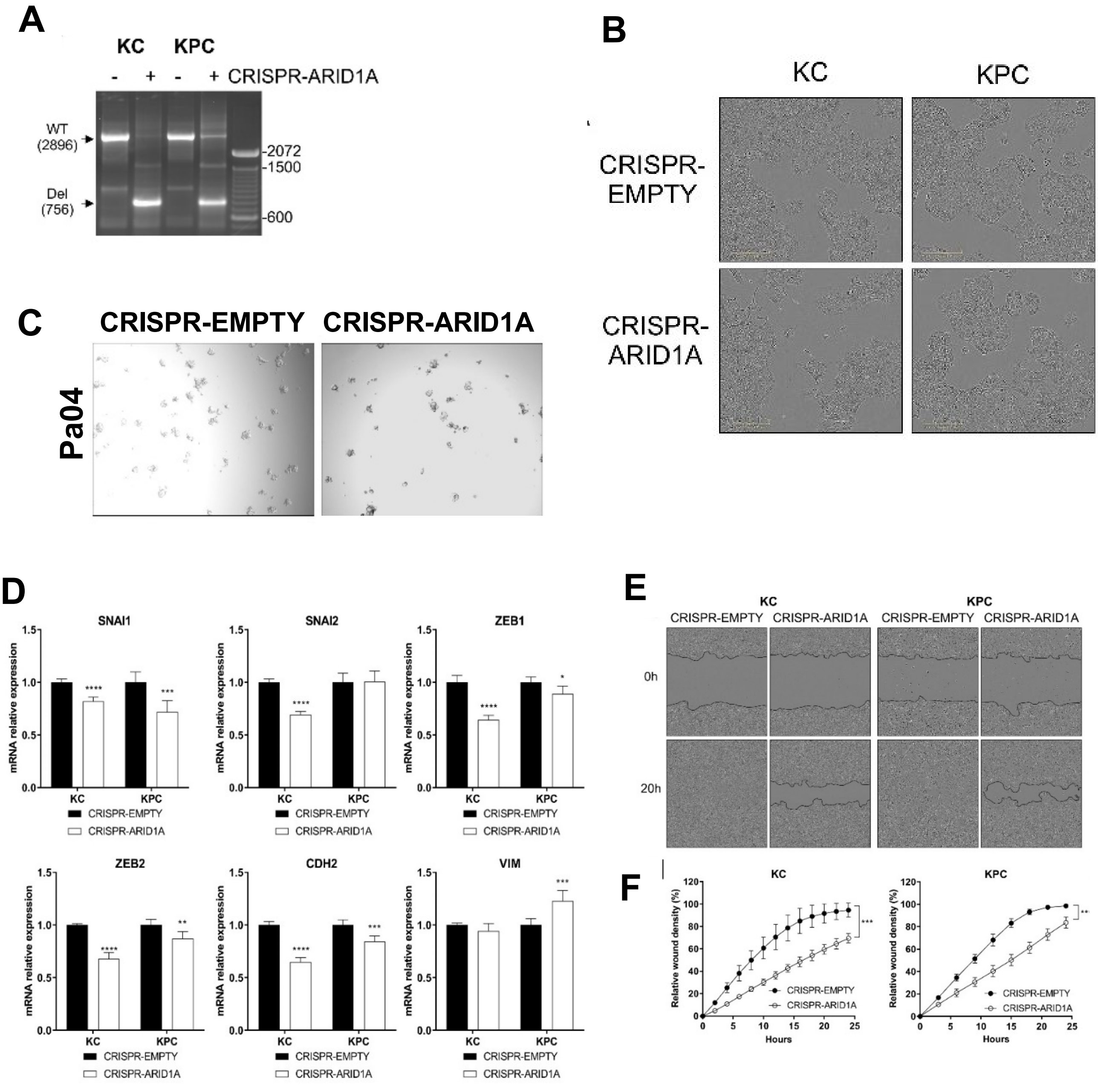


### SUPPLEMENTAL FIGURE S3

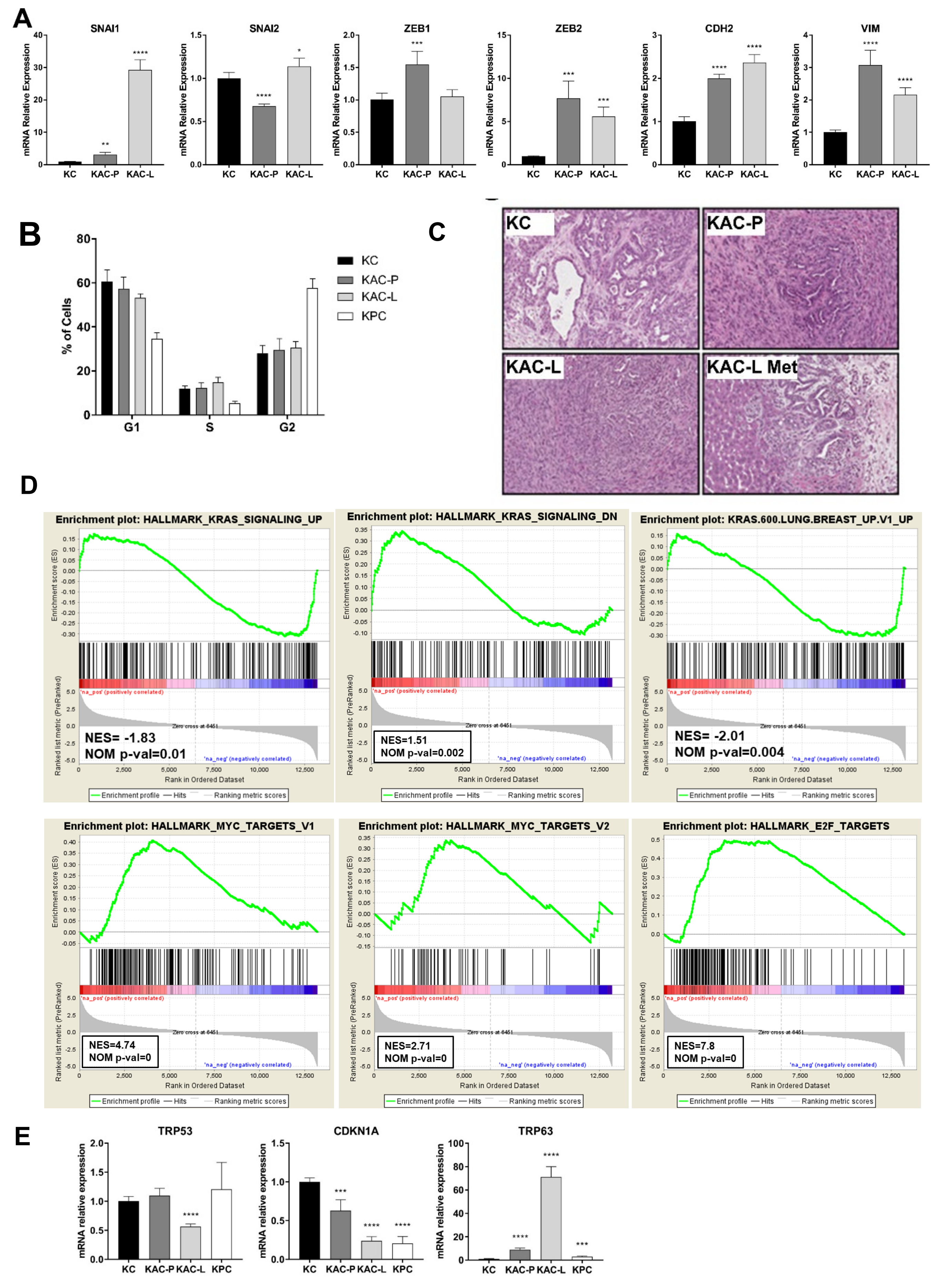


### SUPPLEMENTAL FIGURE S4

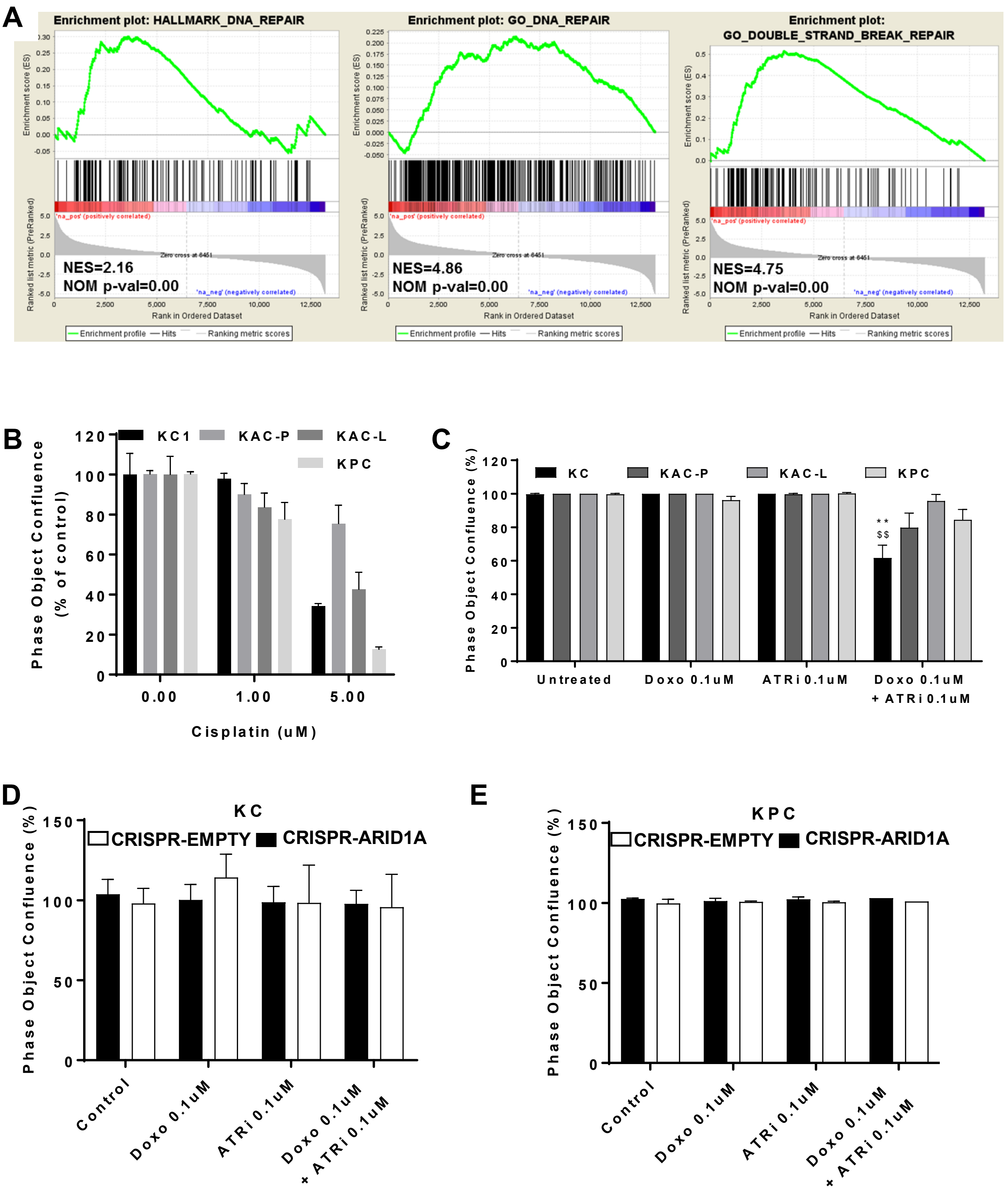


### SUPPLEMENTAL FIGURE S5

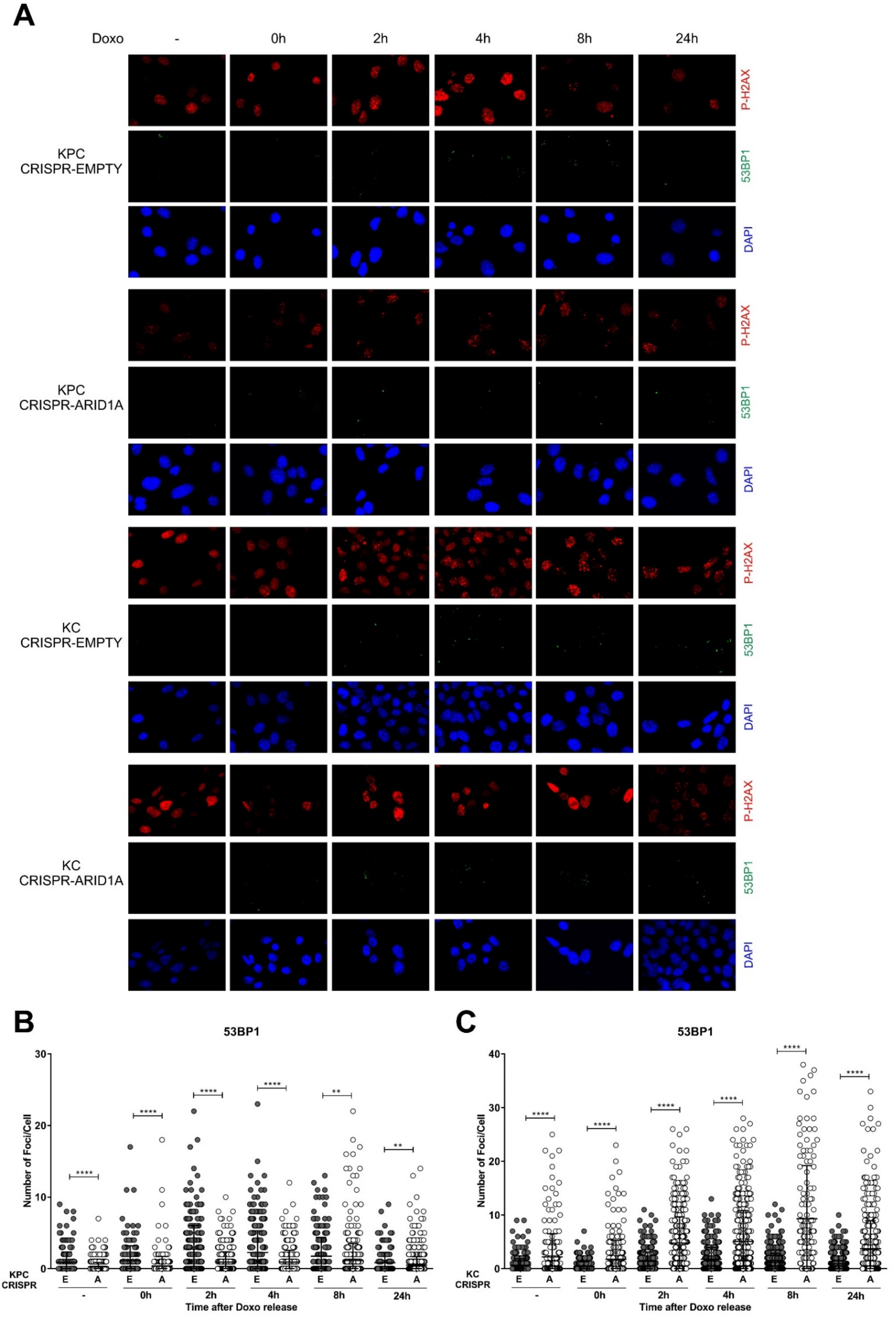


### SUPPLEMENTAL FIGURE S6

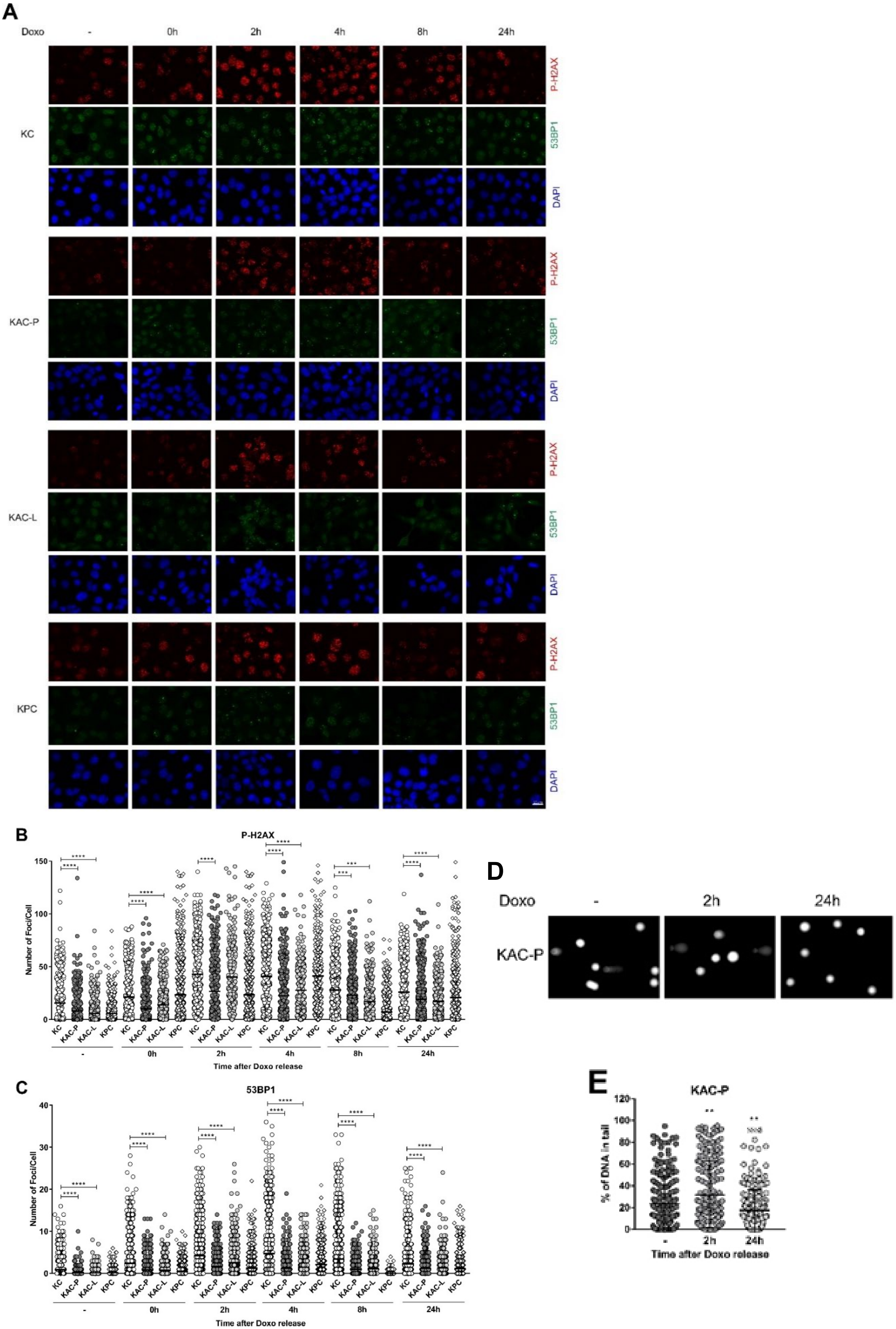


SUPPLEMENTAL FIGURE S7

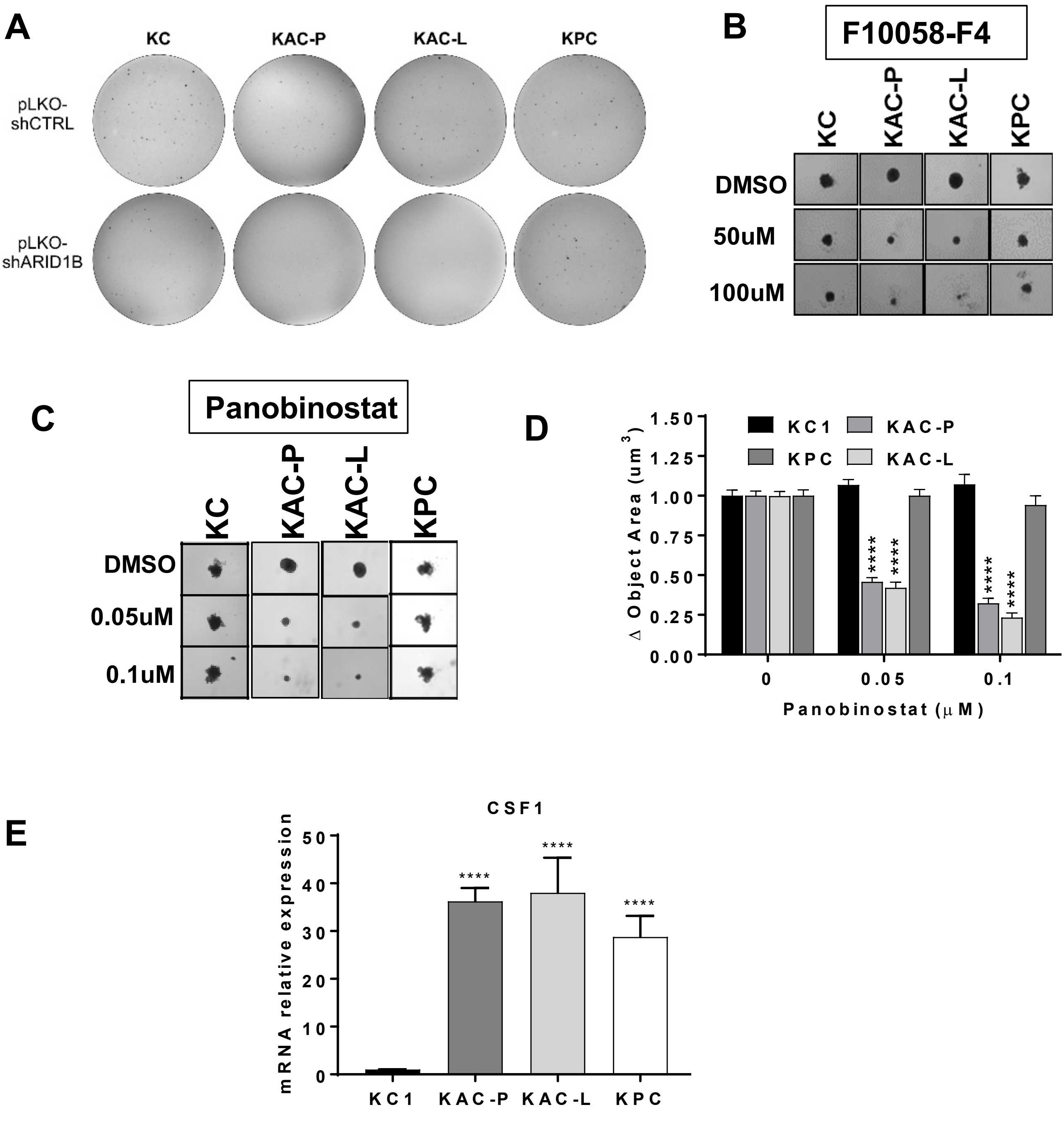


### SUPPLEMENTAL FIGURE S8

#### SUPPLEMENTARY FIGURE LEGENDS

**Supplementary Figure S1. Loss of Arid1a in murine pancreas leads to loss of epithelial homeostasis.** *Arid1a$^{fl/fl}$;Ptf1a-Cre* ("AC") mice were generated by crossing *Ptf1a-Cre* and *Arid1a$^{fl/fl}$* mice, necropsied at regular intervals and pancreas fixed in formalin for histological assessment. **A,** Representative microscopic images of H&E-stained pancreatic sections from "AC" mice at indicated ages revealed widespread parenchymal atrophy accompanied by inflammation starting at 8-weeks, with progressive fatty replacement of normal parenchyma and dilated ducts at 16- and 24-weeks, resembling pancreatitis. *Upper panel*, low magnification at 4x objective lens, *Bottom panel*, high magnification at 4x objective lens. **B,** Gross images of necropsied "AC" mice at indicated older ages (*left panel*) revealed fluid-filled macroscopic cysts, while representative microscopic images of H&E-stained pancreatic sections at low magnification (*middle panels*) and high magnification (*right panels*) revealed complete loss of normal parenchyma accompanied by large dilated mucinous cystic ducts and few LG-PanINs. **C,** Representative microscopic images of IHC on pancreatic sections from 40-wk old "AC" mice confirmed lack of ARID1A expression in Ac: acini; Is: islet; ADM (*left panel*) and Cys: cysts (*right panel*) Scale bar is 100u.

**Supplementary Figure S2. Immunohistochemical characterization of "KAC" pancreas. A,** Representative microscopic images of H&E-stained pancreatic sections from "KAC" mice at indicated ages revealed vast network of mucinous cysts resembling low-grade branched duct gastric type IPMN (LG-IPMN) in humans, admixed with ADM and LG-PanINs. B, Representative microscopic images of IHC for ARID1a in pancreatic sections from "KC" and "KAC" mice at indicated ages, confirmed complete lack of Arid1a expression in "KAC", as expected from a *Ptf1a-Cre* driver line. Representative high-magnification images of IHC staining on pancreata from "KC" (*upper panels*) and "KAC" (*lower panels*) mice for Ki67 **(C)** and cleaved caspase-3 **(D)**. Scale bar is 100u.

**Supplementary Figure S3. Characterization of *Arid1a*-deleted "KC" and "KPC" cell lines using CRISPR-Cas9 system**. **A,** Genomic PCR on DNA isolated from puromycin-selected "KC" and "KPC" cell lines transfected with CRISPR/Cas9 plasmids, detected wild-type (2896 bp) and deleted (756 bp) ARID1A bands. **B,** Microscopic images showing cellular morphology of *Arid1a*-deleted "KC" and "KPC" cell lines. Scale bars, 100 µm. **C,** Representative images of *Arid1a*-deleted Pa04 isogenic cell lines colony growth on soft agar. **D,** Semi-quantitative $RT^2$ PCR for various EMT-associated genes on RNA extracted from *Arid1a*-deleted "KC" and "KPC" isogenic cell lines. Values are expressed as relative expression compared to CRISPR-EMPTY for each cell line. **E,** Representative microscopic images showing migration of *Arid1a*-deleted "KC" and "KPC" cell lines, using Incucyte ZOOM real-time tracking assay. Cells grown to confluence were scratched using the Incucyte Wound maker tool and wound closure tracked every 2 hours post-scratch wound. **F,** Quantification of wound density data represented as mean±SD. Results shown are representative from three independent experiments.

**Supplementary Figure S4. Characterization of autochthonous mouse *Arid1a*-null PDAC cell lines**. **A,** Semi-quantitative $RT^2$ PCR for various EMT-associated genes on RNA extracted from "KC" and "KAC" cell lines revealed higher expression of these genes. Ct values were normalized to Gusb and fold change expression was relative to "KC" group. **B,** Propidium iodide staining was done on cultured cell lines followed by flow cytometric analyses to assess DNA content in various phases of cell cycle. Histogram indicates the percentage of cell lines in each phase of the cell cycle. **C,** Representative microscopic images of H&E-stained sections of mouse pancreatic tumors harvested 4 weeks after orthotopically implanted with "KC", "KAC-P" and "KAC-L" cells in the pancreas of athymic nude mice (N=7 mice per group). "KAC-L" cells also metastasized to liver (KAC-L Met). Scale bar is 100u. **D,** Gene set enrichment analysis (GSEA) of differentially expressed transcripts in RNA-Seq of "KC" and "KAC" cells showed enrichment in gene sets associated with downregulated Kras signaling (*upper panel*) and positive enrichment in hallmark gene signature for Myc and E2F targets (*lower panel*). **E,** Semi-quantitative $RT^2$ PCR showing reduced expression of direct p53 target gene CDKN1A and increased expression of Trp63 in “KAC” cell lines, relative to “KC”. “KPC” cells with mutant p53 also showed reduced CDKN1A expression, as expected. Ct values were normalized to Actin and fold change expression was relative to “KC” group. Representative findings from at least 3 independent experiments are shown and data analyzed using the two-tailed unpaired Student’s t test and considered significant if *, $P< 0.05$; **, $P< 0.01$; ***, $P< 0.001$; ****, $P< 0.0001$, unless otherwise specified.

**Supplementary Figure S5. PDAC cells from *Arid1a*-null “KAC” mice shows enhanced DNA repair capability**. **A,** GSEA analysis of differentially expressed transcripts in RNA-Seq of “KC” and “KAC” cells showed enrichment in signatures associated with DNA repair in “KAC” cells compared to KC control. **B,** Monolayer culture of autochthonous PDAC cell lines were treated either with increasing doses of Cisplatin (B) or combination of Doxorubicin and ATRi (C) for 72h. Cell confluency was imaged and quantified by Incucyte ZOOM and data represented as % of the corresponding vehicle control for each cell line. ****, $P< 0.0001$ for 5uM Cisplatin. **D-E,** Sensitivity of *Arid1a*-deleted isogenic “KC” (D) and “KPC” (E) cell lines to 72h of Doxorubicin and ATRi treatment, evaluated as in (B). Representative findings from at least 3 independent experiments are shown and data analyzed using the two-tailed unpaired Student’s t test. **, $P< 0.01$ when compared to Doxo alone; $^{\$\$}$, $P< 0.01$ when compared to ATRi alone.

**Supplementary Figure S6**. **Assessment of DNA damage post *Arid1a*-deletion in PDAC cells**. **A,** Immunofluorescence staining for P-H2AX, 53BP1 and DAPI showing nuclear foci after Doxo treatment in *Arid1a*-deleted isogenic “KC” and “KPC” cell lines. Cells were exposed to 0.1uM Doxo for 30min and fixed at indicated timepoints. Representative images shown from at least 3 independent experiments. Scale bar, 20 μm. **B-C,** Scatter plots showing quantification of the number of 53BP1 foci/cell in “KPC” (left) or “KC” (right) isogenic cell lines, performed with iMaris

Microscopy Image Analysis Software (Bitplane). Data are represented as mean±SD and two-tailed unpaired Student's t test have been used for data analysis (unless otherwise indicated) and considered significant if *, *P*< 0.05; **, *P*< 0.01; ***, *P*< 0.001; ****, *P*< 0.0001, unless otherwise specified.

**Supplementary Figure S7**. **Assessment of DNA damage repair in autochthonous mouse PDAC cell lines. A,** Immunofluorescence staining for P-H2AX, 53BP1 and DAPI showing nuclear foci after Doxo treatment in autochthonous *Arid1a*-null cell lines. Cells were exposed to 0.1uM Doxo for 30min and fixed at indicated timepoints. Representative images shown from at least 3 independent experiments. Scale bar, 20 µm. **B-C,** Scatter plots showing quantification of the number of P-H2AX (B) or 53BP1 (C) foci/cell, performed with iMaris Microscopy Image Analysis Software (Bitplane). **D-E,** Representative images from Comet assay performed after exposing KAC-P cells to 0.1uM Doxo for 30 min and then released for 2 or 24h. Quantification indicated as mean±SD with two-tailed unpaired Student's t test used for data analysis; *, *P*< 0.05; **, *P*< 0.01; ***, *P*< 0.001; ****, *P*< 0.0001. *two-tailed unpaired Student's t test against untreated sample, $two-tailed unpaired Student's t test against 2h timepoint.

**Supplementary Figure S8**. **Synthetic lethality in autochthonous *Arid1a*-null PDAC cells. A,** Representative well images of colonies in soft agar, with 14d of growth post transduction with pLKO-shARID1B vector. **B,** Representative images of F10058-4 treated spheroids from autochthonous cell lines grown on ultralow attachment plates for 7d. **C-D,** Representative images of Panobinostat treated spheroids from autochthonous cell lines grown on ultralow attachment plates for 7d. Images were captured and spheroid Area (um3) measured using spheroid imaging protocol of the Gen5 Image software on Cytation 3 (Biotek) using 10X objective lens; data normalized to vehicle control and plotted as change relative to control. **E,** Relative expression levels of Csf1 assessed by semi-quantitative $RT^2$ PCR in "KAC" cell lines. Ct values were normalized to Actin and fold change expression was relative to "KC" group. Two-tailed unpaired

Student's t test have been used for data analysis and considered significant if *, $P < 0.05$; **, $P < 0.01$; ***, $P < 0.001$; ****, $P < 0.0001$.

#### Supplementary Material and Methods

##### Orthotopic Implantation

Orthotopic tumors were established in athymic nude mice, as per established protocol (47). Briefly, cells were suspended in ice-cold PBS and Matrigel in a 1:1 ratio, and injected into the pancreatic parenchyma of anesthetized mice.

##### Viral Transduction

Lentiviral infections were performed using 293LTV cells (Cell Biolabs, Cat#LTV-100) as producers of viral supernatants, upon co-transfection with MISSION shRNA lentiviral DNA (Sigma-Aldrich) and the helper vectors pCMVR8.74 (Addgene Cat#22036) and pCMV-VSV-G (Addgene, Cat #8454) using Lipofectamine 3000 transfection reagent (ThermoFisher Scientific, Cat# L3000008). Supernatants from 293LTV cultures was passed through a 0.45 µm filter before transduction of cancer cells, with successful clones selected by 1.5µg/mL puromycin (InvivoGen, Cat# ant-pr1). Sequence of ARID1B shRNA (TRCN0000238628) used was as follows: CCGGGCCGAATTACAAACGTCATATCTCGAGATATGACGTTTGTAATTCGGCTTTTTG. The control scrambled shRNA (Sigma-Aldrich, Cat# SHC002) does not target any known mouse gene.

##### Antibodies and reagents

Antibodies used are as follows: Anti-ARID1A (Cat#12354), anti-ACTIN (Cat#4970), anti-P44/42 MAPK (Cat#4695), anti-pP44/42 MAPK(Erk1/2)(Thr202/Tyr204) (Cat#4370), anti-p53 (rodent specific, Cat#32532), anti-MSH2 (Cat#2017), anti-MSH6 (Cat#3995), anti-SOX2 (Cat#14962), anti-Phospho-H2A.X(Ser139) (for WB, Cat#9718), anti-Mouse HRP-linked (Cat#7076), anti-Rabbit HRP-linked (Cat#7074) from CST; Anti-p21 (Cat#ab1019199) and anti-MLH1 (Cat#ab92312) from Abcam; Anti-Phospho-H2A.X (Ser139) (Cat#05-636) from Millipore; Anti-ARID1B (Cat#A301-046A) from Bethyl Laboratories; Anti-Claudin 18 (Cat#7000178), anti-Rabbit Alexa Fluor 488 (Cat#A21206), anti-Mouse Alexa Fluor Plus 555 (Cat#A32727) from ThermoFisher Scientific; Anti-PMS2 (Cat#556415) and PI/RNase staining solution (Cat#550825) from BD Biosciences; Anti-53BP1 (Cat#NB100-304) from Novus Biologicals. GSK126, F10058-F4, Panobinostat, ATRi (VE822), Cisplatin and Doxorubicin, were purchased from Selleckchem.

**Flow Cytometric Analyses**

Cell cycle profiles were measured by flow cytometry using propidium iodide (PI) as described before [13].

**Western blot analysis**

Immunoblotting was performed as described before [13]. Briefly, cellular proteins were extracted in RIPA lysis buffer (Sigma-Aldrich, Cat#R0278) plus protease and phosphatase inhibitors (Sigma-Aldrich, Cat#P0044, P5726, P8340) and separated by SDS–PAGE. Membranes with transferred protein were incubated with primary antibodies at 4°C overnight followed by HRP-conjugated secondary antibodies and visualized using the enhanced chemiluminescence (ECL) detection system (Biorad, Cat#1705061) by ChemiDoc (Biorad) scanning.

**Quantitative RT–PCR**

Total RNA was extracted using the RNeasy Mini kit (Qiagen, Cat#74106) and reverse transcription performed using High-Capacity cDNA Reverse Transcription Kit (ThermoFisher Scientific, Cat#4368813) according to the manufacturer's instructions. Quantitative RT–PCR were performed using the StepOnePlus Real-Time PCR System (ThermoFisher Scientific), with predesigned TaqMan gene expression assays (ThermoFisher Scientific): Actb (Mm00607939_s1), ARID1B (Mm01338353_m1), SNAI1 (Mm00441533_g1), SNAI2 (Mm00441531_m1), ZEB1 (Mm00495564_m1), ZEB2 (Mm00497196_m1), VIM (Mm01333430_m1), CDH2 (Mm01162497_m1), SOX2 (Mm03053810_s1), CLDN18 (Mm00517321_m1) and EZH2 (Mm00468464_m1). RT-PCR reactions were performed in triplicate and analysis done as described before[13] with data normalized on "KC" as control.

**MMR assay**

The MMR assay was performed as previously described [14]. Briefly, cells were seeded in 12-well plates and were transfected with a plasmid mixture containing 160 ng of pmax-mOrange (vector control) or pmax-G:G-mismatch-mOrange (MMR) together with 840 ng of carrier DNA . After 48 hours of transfection, cells were harvested and analysed by BD FACSCELESTA flow cytometer (BD Biosciences). The relative MMR capacity, expressed as % of reporter expression (R.E.) was determined by dividing the percentage of mOrange-positive cells in MMR by the percentage of mOrange-positive cells in vector control.

**RNA-Sequencing and Bioinformatic analysis**

Total RNA was isolated from cultured "KC" and "KAC" cell lines, using RNeasy Mini kit (Qiagen, Cat#74106) and integrity validated using the Agilent 2100 Bioanalyzer. Libraries were generated using 1–2μg of total RNA following the Illumina protocol for preparing samples for high throughput stranded mRNA sequencing on the Illumina NextSeq 500 High sequencer using the 76nt PE format. As described previously[1], sequence reads were aligned to the GRCm38 mouse genome with tophat (v2.0.13) with parameters allowing a read to be mapped to at most one location: "--no-coverage-search -p 1 -g 1". Gene hits were counted with HTseq (v0.6.1) under default parameters using release 84 of the Mus_musculus.GRCm38 GTF annotation file. Differential expression analysis was performed in R/Bioconductor following the DESeq2 workflow [2] and annotated with biomaRt.

**ATAC-seq**

A suspension of 50,000 cells from both "KC" and "KAC" cultures in cold lysis buffer was incubated in transposition reaction mixture at 37°C and subjected to ATAC-seq, as described previously[1]. Briefly, after the reaction was immediately purified by the Qiagen MinElute PCR Purification Kit, PCR was performed on the eluted DNA using barcoded primer and thermal cycle according to the method described above. We used Bowtie 2 (version 2.2.3) with parameters allowing for soft clipping to align the sequencing reads for each sample to the NCBI reference mouse genome sequence (GRCm38). Peak calling for each sample was performed using MACS2 (version 2.1.0) with default parameters. For discovering differential peaks between samples, a new set of peak ranges was constructed, formed from the union of individual peaks, merging those within 200bp of each other, and then enumerated by counting the reads mapped to each region. Peak region counts were normalized following the DESEQ2 workflow, annotated with biomaRt, and genes were selected by a threshold on fold change. Identification of transcription factor binding site motifs was done using HOMER with default parameters for motif identification[3].

**Supplementary Table 1.** **Immunohistochemical staining for ARID1A on human IPMNs classified based on dysplasia grade and subtypes**

| Sample# | ARID1A Expression | Dysplasia grade | IPMN Subtype |
|---|---|---|---|
| IPMN-1 | - | Low | Gastric |
| IPMN-2 | - | Low | Gastric |
| IPMN-3 | - | Low | Gastric |
| IPMN-4 | - | Low | Gastric |
| IPMN-5 | - | Low | Gastric |
| IPMN-6 | - | Low | Gastric |
| IPMN-7 | - | Low | Gastric |
| IPMN-8 | - | Low | Gastric |
| IPMN-9* | - | Low | Gastric |
| IPMN-9* | +++ | High | Gastric |
| IPMN-10* | - | Low | Gastric |
| IPMN-10* | +++ | High | Gastric |
| IPMN-11 | + | Low | Gastric |
| IPMN-12 | + | Low | Pancreatobiliary |
| IPMN-13 | + | Low | Gastric |
| IPMN-14 | + | Low | Gastric |
| IPMN-15 | + | Low | Gastric |
| IPMN-16 | + | Low | Gastric |
| IPMN-17 | + | Low | Gastric |
| IPMN-18 | + | Low | Gastric |
| IPMN-19 | + | Low | Gastric |
| IPMN-20 | + | Low | Gastric |
| IPMN-21 | + | Low | Pancreatobiliary |
| IPMN-22 | + | Low | Gastric |
| IPMN-23 | + | Low | Gastric |
| IPMN-24 | + | Low | Gastric |
| IPMN-25 | + | Low | Gastric |
| IPMN-26 | + | Low | Gastric |
| IPMN-27 | + | Low | Gastric |
| IPMN-28 | + | Low | Gastric |
| IPMN-29 | + | Mix | Pancreatobiliary |
| IPMN-30 | + | Mix | Pancreatobiliary |
| IPMN-31 | + | Mix | Pancreatobiliary |
| IPMN-32 | + | Mix | Gastric |
| IPMN-33 | + | Mix | Gastric |
| IPMN-34 | + | Mix | Intestinal |
| IPMN-35 | + | High | Intestinal |
| IPMN-36 | + | High | Gastric |
| IPMN-37 | + | High | Gastric |
| IPMN-38 | + | High | Pancreatobiliary |
| IPMN-39 | + | High | Pancreatobiliary |
| IPMN-40 | + | High | Pancreatobiliary |
| IPMN-41 | + | High | Gastric |
| IPMN-42 | + | High | Gastric |
| IPMN-43 | + | High | Pancreatobiliary |
| IPMN-44 | + | High | Gastric |
| IPMN-45 | + | High | Intestinal |
| IPMN-46 | + | High | Intestinal |
| IPMN-47 | + | High | Intestinal |
| IPMN-48 | + | High | Gastric |
| IPMN-49 | ++ | High | Pancreatobiliary |
| IPMN-50 | ++ | High | Pancreatobiliary |
| IPMN-51 | ++ | High | Pancreatobiliary |
| IPMN-52 | ++ | High | Intestinal |
| IPMN-53 | ++ | High | Intestinal |

*Cases with regions of both low and high grade dysplasia
